# Stiffness Sensing by Smooth Muscle Cells: Continuum Mechanics Modeling of the Acto-Myosin Role

**DOI:** 10.1101/2023.05.02.539179

**Authors:** Ali Akbar Karkhaneh Yousefi, Claudie Petit, Amira Ben Hassine, Stéphane Avril

**Author notes:** Corresponding Author: Stéphane Avril.

## Abstract

Aortic Smooth Muscle Cells (SMCs) play a vital role in maintaining homeostasis in the aorta by sensing and responding to mechanical stimuli. However, the mechanisms that underlie the ability of SMCs to sense and respond to stiffness change in their environment are still partially unclear. In this study, we focus on the role of acto-myosin contractility in stiffness sensing and introduce a novel continuum mechanics approach based on the principles of thermal strains. Each stress fiber satisfies a universal stress-strain relationship driven by a Young’s modulus, a contraction coefficient scaling the fictitious thermal strain, a maximum contraction stress and a softening parameter describing the sliding effects between actin and myosin filaments. To account for the inherent variability of cellular responses, large populations of SMCs are modeled with the finite-element method, each cell having a random number and a random arrangement of stress fibers. Moreover, the level of myosin activation in each stress fiber satisfies a Weibull probability density function. Model predictions are compared to traction force measurements on different SMC lineages. It is demonstrated that the model not only predicts well the effects of substrate stiffness on cellular traction, but it can also successfully approximate the statistical variations of cellular tractions induced by intercellular variability. Finally, stresses in the nuclear envelope and in the nucleus are computed with the model, showing that the variations of cytoskeletal forces induced by substrate stiffness directly induce deformations of the nucleus which can potentially alter gene expression. The predictability of the model combined to its relative simplicity are promising assets for further investigation of stiffness sensing in 3D environments. Eventually, this could contribute to decipher the effects of mechanosensitivity impairment, which are known to be at the root of aortic aneurysms.

## 1 Introduction

There is growing evidence that stiffness increase is a sign of weakness in many soft tissues, including the aortic wall.^1–3^ In Ascending Thoracic Aortic Aneurysm (ATAA) for instance, which is one of the most serious aortic diseases that can lead to catastrophic complications,^4, 5^ stiffness increase is even a significant factor for the risk of dissections.^6, 7^

In the healthy aorta, Smooth Muscle Cells (SMCs) are essential for the regulation of the wall stiffness.^8–16^ Thanks to their phenotypic plasticity, SMCs can maintain mechanical homeostasis through variations of their active tone (contractile phenotype, short-term adaptation) and through synthesis and remodeling of the extracellular matrix (ECM) (synthetic phenotype, long-term adaptation).^1, 2, 14, 17–20^ However, missensing of mechanical stimuli by SMCs (stress, strain, or stiffness) can alter the maintenance of mechanical homeostasis and induce impaired adaptations that are responsible for ATAA progression, for instance in the Marfan syndrome.^2, 21–24^ Accordingly, there is a pressing need to better investigate and model SMC biomechanics and its sensitivity to ECM stiffness.

However, the mechanisms that underlie the ability of SMCs to sense and respond to stiffness change are still partially unclear. Several previous studies on different cell types have examined the roles of mechanoreceptors at the cellular membrane.^25–27^ Moreover, recent studies have highlighted the significant role of contractile acto-myosin units in the cytoskeleton that can act as stiffness sensors.^28–30^ In the current study, we investigate stiffness sensing through the contractile acto-myosin units.

To the best of the authors’ knowledge, only a small number of studies investigated the relationship between the contractility of aortic SMCs and ECM stiffness.^31^ Recently, Petit and colleagues quantified the basal tone of SMCs at the single-cell level by conducting Traction Force Microscopy (TFM) tests on aortic SMCs cultured on gels of different stiffness properties.^32, 33^ They reported that the traction forces of SMCs significantly depend on the elastic modulus of the gels on which they are cultured. Moreover, vascular SMCs showed a highly anisotropic behavior^34^ and an intrinsic ability to modulate the load borne by the surrounding ECM.^35^

The main contributors to the overall mechanical behavior of SMCs at the single-cell level are the cell membrane, the cytoplasm, stress fibers, the nucleus, and the nuclear envelope. The latter should also include a perinuclear actin cap connecting a fraction of stress fibers to the interphase nucleus through linkers of nucleoskeleton and cytoskeleton (LINC) protein complexes.^36, 37^ A number of computational models in cell biomechanics did not take into account all these components, but regarded the whole cell as a black box with global mechanical properties to predict the overall passive response.^34, 35, 38^ Other computational models were proposed to include cell contraction.^29, 39–43^ These models do not always account for cell mechanosensitivity^40^ and those relating cellular contraction to the environmental stiffness, such as the motor-clutch-based models, often require a large number of constitutive parameters.^39, 41^ Moreover, none of these models has ever attempted to consider the effects of the cell geometry or of the arrangements of stress fibers within finite-element (FE) analyses.

The main goal of this study is to develop a new computational model for the contractile behavior of aortic SMCs (ASMCs) that can address those shortcomings and account for the variations of cellular traction with the surrounding stiffness. In section 2, we describe the new model of ASMCs and the different experimental arrangements to conduct TFM analyses on two different ASMC lineages. In section 3, the proposed model is calibrated against the TFM results and used to evaluate the effects of substrate stiffness on the stresses in the nuclear envelope and the nucleus. Section 4 is dedicated to the discussion of results.

## 2 Materials and Methods

### 2.1 Computational model of ASMCs

#### 2.1.1 Mechanical model of stress fibers

Stress fibers represent cytoskeletal truss-like structures composed of cross-linked actin filament bundles and myosin motor proteins. Myosin activation, stimulated by neurotransmitters, hormones, and ionic channel, results in stress fiber shortening, whereas deactivation results back into stress fiber extension.

Recent findings showed that the myosin filaments contract the actin filaments to a fixed distance.^44^ Inspired by this, in this study, we assume that stress fibers uniformly shorten after myosin activation. As ASMCs are attached to the substrate at focal adhesions, shortening results in a deformation of the substrate. If the substrate is sufficiently compliant, the deformation is large and the stress fibers only withstand a small tensile force, but if the substrate is rigid, the deformation becomes negligible and the stress fibers have to withstand large tensile forces. Therefore, traction forces tend to increase with the substrate stiffness, which is consistent with recent TFM results on ASMCs.^39, 45^ However, when the substrate becomes too stiff, traction forces can potentially reach a certain elastic limit, which may correspond to sliding between actin and myosin filaments.^39, 43^ As predicted by the motor–clutch-based model, there exist an ‘optimal stiffness’ where cells can generate maximal traction.^46^

To model shortening of stress fibers induced by myosin activation, we introduced an eigenstrain, denoted *α*Δ*T*, analogously to a thermal contraction:

- *α* is the contraction coefficient, which relates the level of stress fiber shortening to the level of myosin activation. It is a dimensionless strictly positive parameter.
- Δ*T* represents the level of myosin activation, satisfying 0 ≤ Δ*T* ≤ 1, in which Δ*T* = 1 corresponds to the maximal activation and Δ*T* = 0 to no activation (full relaxation).^42^ In the normal or basal conditions investigated by the current study, the activation level is assumed to satisfy a Weibull probability density function, which may be written:

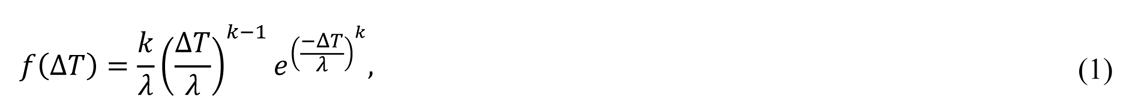

where *k* > 0 represents the shape parameter of the distribution (*k*=1 for the exponential distribution, *k*=2 for the Rayleigh distribution) and *λ* > 0 is the scale parameter of the distribution. It is related to the median of the distribution, denoted 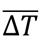, according to:

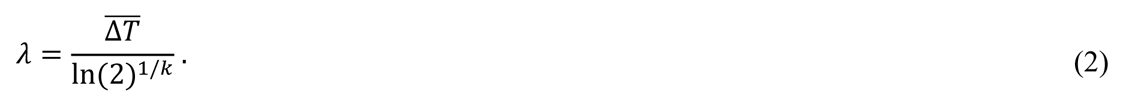

For instance, *λ* ≈ 1.44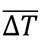 for the exponential distribution (*k*=1) and *λ* ≈ 1.2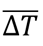 for the Rayleigh distribution (*k*=2). The Weibull distribution law is not literally bounded on the [0,1] interval but *f*(Δ*T*) always took very low values (below 0.0005) for Δ*T*=1 in the different distributions measured during this study, making the Weibull function well suited to represent the distributions of experimental myosin activations on the [0,1] interval. Along with the varying number of stress fibers between cells, the variations of Δ*T* is a major source of intercellular variations among the traction forces applied by ASMCs on their substrate.

Let us consider an isolated fictitious SMC where Δ*T* = 0. Each stress fiber has an initial length *L*_0_ in these conditions. The *α*Δ*T* eigenstrain refers to the shortening of stress fibers when the same isolated SMC is subject to some myosin activation Δ*T* > 0 (basal tone in our case here). The obtained fictitious length of stress fibers would then be: *L* = (1 − *α*Δ*T*)*L*_0_. However, as the SMC is not isolated but adheres to a substrate, the actual shortening of stress fibers satisfies *ε*_*a*_ < *α*Δ*T*, and the actual length of stress fibers is then *L* = (1 − *ε*_*a*_)*L*_0_.

To model numerically the mechanical behavior of stress fibers, we assumed that stress fibers have a stress-strain response that is composed of two parts (Figure 1):

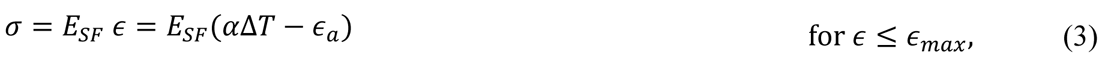

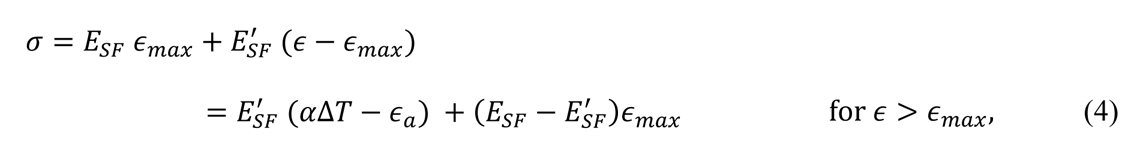

in which *σ* and *ε* represent the nominal stress and the elastic nominal strain of a truss element modeling the stress fiber, *E*_*SF*_is the Young’s modulus and *σ*^′^ is the softening coefficient. Softening for *ε* > *E*^’^_*SF*_is attributed to the sliding effects between actin and myosin filaments beyond the maximum contraction stress. It is known that filament overlap between actin and myosin decreases beyond a certain stretch level, inducing a decrease of the active stress.^43^ This is also in agreement with the motor–clutch-based model which predicts the existence of an ‘optimal stiffness’ where cells can generate maximal traction.^46^

**Figure 1.**
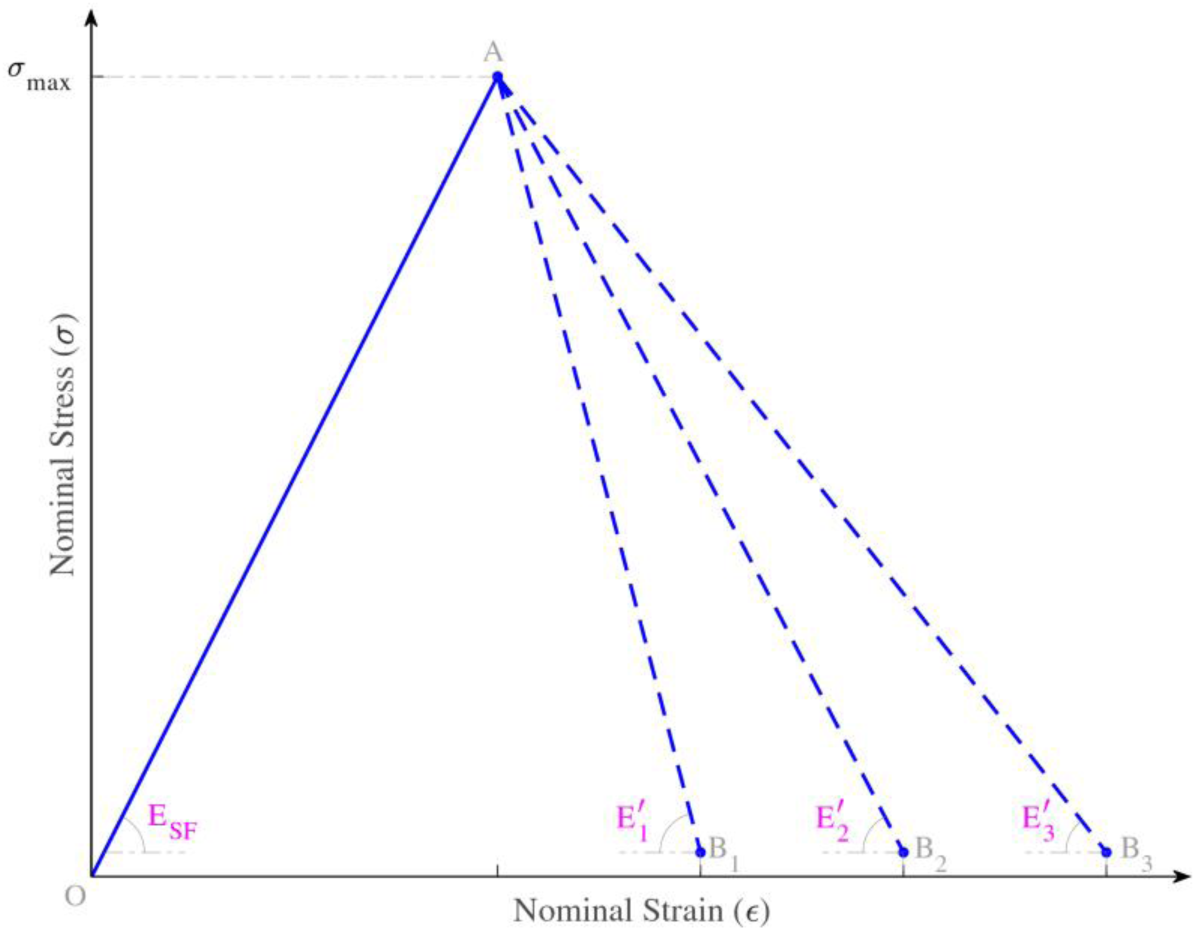
Schematic stress-strain behavior of a stress fiber. The stress can increase with the strain over OA with a slope denoted *E*_*SF*_(Young’s modulus of stress fibers), until reaching a critical stress value denoted *σ*_*max*_(elastic limit). Beyond *σ*_*max*_, the stress decreases with a slope denoted *E*^′^. This softening effect is attributed to the sliding effects between actin and myosin filaments.

As stress fibers with a higher myosin activation also have a higher capacity to withstand traction forces due to a larger number of active bridges between actin and myosin,^43^ it was assumed that the strain threshold satisfies: *ε*_*max*_ = *ξα*Δ*T*, where *ξ* is a linear coefficient such as 0 < *ξ* < 1.

In summary, five constitutive parameters had to be adjusted in our stress fiber model:

1. the Young’s modulus *E*_*SF*_,
2. the level of myosin activation Δ*T*,
3. the contraction coefficient *α*,
4. the linear coefficient *ξ* defining the elastic limit *ε*_*max*_ = *ξα*Δ*T*,
5. the softening coefficient *σ*^′^ (or *σ*^′^ (*i* = 1,2,3) in Figure 1).

We assumed that the stiffness of a stress fiber was a constant, with the Young’s modulus set to *σ*_*SF*_ = 50 MPa and the cross-section being circular with a diameter 0.2 µm.^47^ The level of myosin activation satisfied a Weibull probability density function driven by 2 parameters which were determined for each cell lineage: *k* and 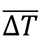. Along with the three other model parameters *α*, *ξ* and *σ*^′^, which were also assumed to vary between the different cell lineages, they were identified with TFM experiments that are presented in subsection 2.2.

#### 2.1.2 FE model of SMCs

As shown in Figure 2, aortic SMCs usually exhibit a spindle shape with focal adhesions at both ends. In our model, we consider a semi-spindle of dimensions 250 x 30 µm^2^ with a cell membrane surrounding this spindle (Figure 2). The cell membrane is directly in contact with the cytoplasm. We also considered a nuclear envelope covering the nucleus and surrounded by the cytoplasm.

**Figure 2.**
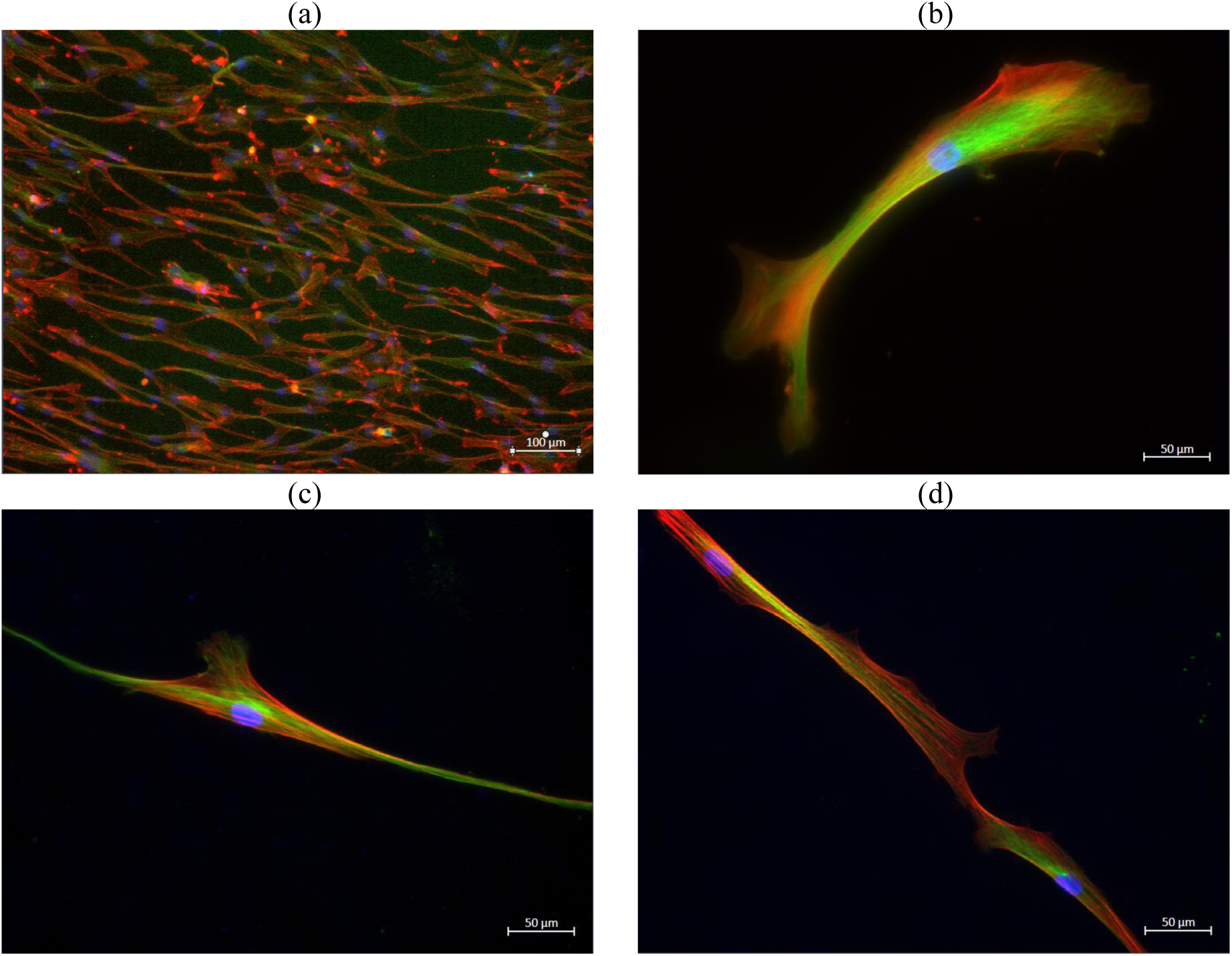
Examples of cultured aortic smooth muscle cells (AoSMC, Lonza) stained with fluorescent markers and observed with fluorescence microscopy. F-actin is stained in red (Phalloidin-Rhodamin fluorescent marker), myosin is stained in green (Alexafluor fluorescent marker) and the nucleus is stained in blue (Hoescht fluorescent biomarker). An AoSMC culture with a high cell density is shown in (a), whereas single AoSMCs are shown in (b), (c) and (d), obtained from cultures with smaller cell densities.

We assumed that the cytoplasm of ASMC includes a random number of stress fibers, this number following a uniform distribution (mean = 25, standard deviation = 6.05) based on experimental observations as shown in Figure 2. Also based on experimental observations shown in Figure 2, we connected at least half of the stress fibers to focal adhesions on both sides, the others being connected to the nuclear envelope at one end. The latter connections are supposed to model the effects of the perinuclear actin cap connecting a fraction of stress fibers to the interphase nucleus through LINC protein complexes.^36, 37^ Schematic views of the ASMC model are shown in Figure 3.

**Figure 3.**
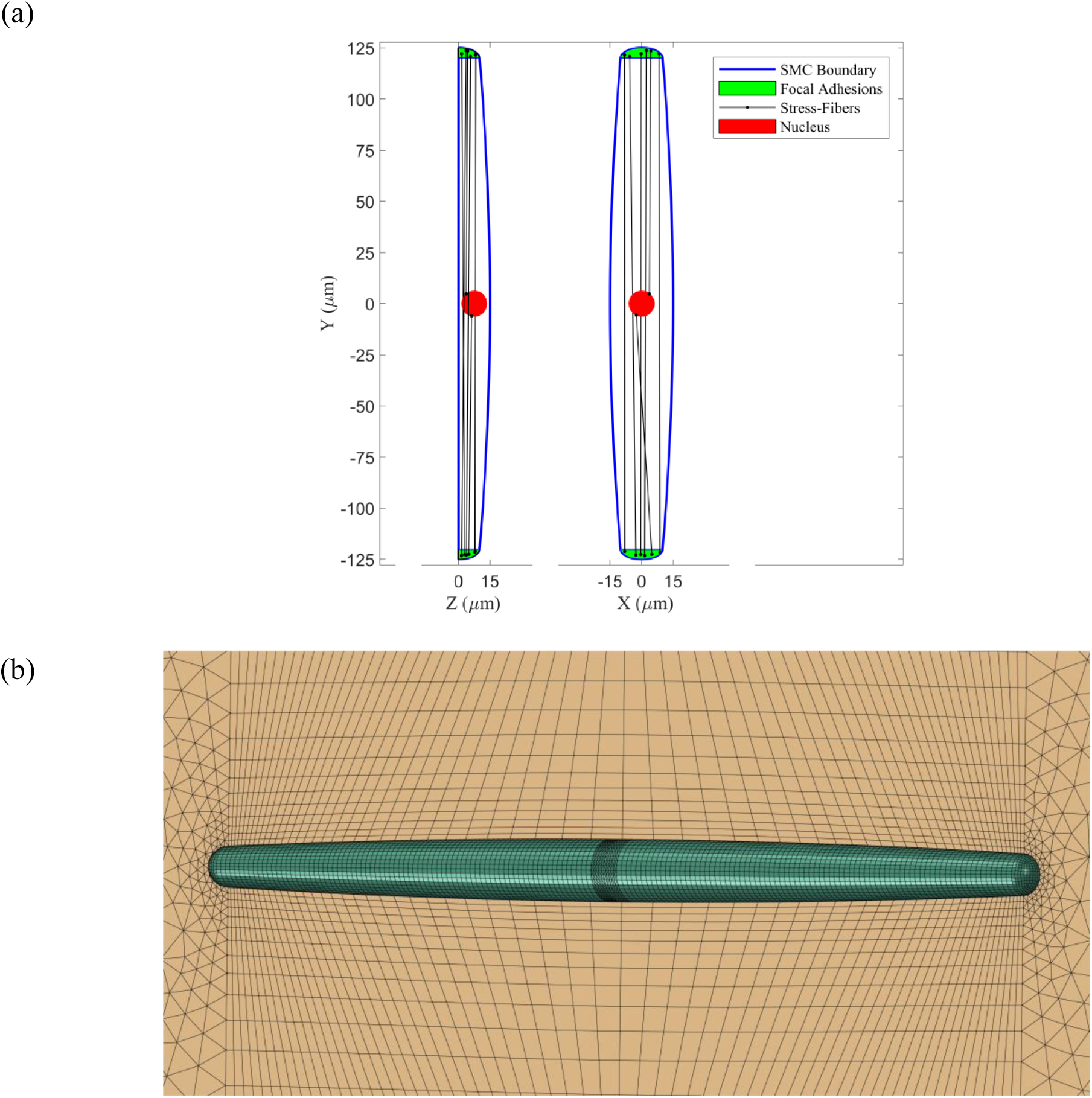
(a): Schematic views of a single SMC on the XY and ZY planes; (b): A 3-D view of the generated mesh on the cell and substrate for FE simulations.

The single 250-µm-long ASMC shown in Figure 3 and the underlying substrate (0.4×5×5 mm^3^) were meshed and computed with the Abaqus software. The cytoplasm was meshed with 16388 solid elements (C3D8 and C3D4), the nucleus with 1024 C3D8 elements, whereas the cell membrane and the nuclear envelope were meshed with 3860 and 192 shell elements (S3 and S4) respectively, both with 10 nm thickness.^48^ The substrate was meshed with 53662 solid elements (C3D8 and C3D4).

A tie constraint was assigned between the nodes of the cell membrane and the nodes of the substrate surface at the focal adhesion (surface of 148.84 µm² corresponding to 171 nodes). We assumed that the sum of reaction forces at all the tied nodes of a focal adhesion (resultant adhesion force) was a prediction of the traction forces measured in TFM experiments.

The whole model was formulated to handle finite deformations. The cell membrane and the nuclear membrane were modeled as neo-Hookean materials with a shear modulus of *C*_10_ = 600 kPa.^49^ A neo-Hookean behavior was also assumed for both the cytoplasm and the nucleus with a shear modulus of *C*_10_ = 100 Pa.^50^ Equivalent Poisson’s ratios of *v* = 0.45 and *v* = 0.499 were assigned to the substrate and to all the cell components, respectively.

#### 2.2 Experimental Measurements

##### 2.2.1 Cell Lineages

For the calibration of our computation model, we used cell cultures of the two following human ASMC lineages:

1. the first cell lineage (named ASMC_1 onwards) is a commercial immortalized human ASMCs lineage purchased from Lonza and delivered at passage 3, obtained from a 30-year-old female donor. The cells were first cultured for initial proliferation in growth medium (SmGM- 2, Lonza). Then, they were frozen into 1.5 mL aliquots containing 10% Fetal Bovine Serum (FBS), 10% Dimethyl sulfoxide as a cryoprotectant, and 80% SmGM-2 complete medium. Each aliquot contained around 3 million cells. The ASMCs were stored into liquid nitrogen at passage 5 for further experiments.
2. the second lineage (named ASMC_2 onwards) is a primary culture from our laboratory. ASMCs were extracted from ATAA tissues collected after informed consent during elective surgical aneurysm repair. This lineage was collected from a 72-year-old male patient. The tissue was stored in physiological serum and put into the incubator at 37°C within two hours after surgery. Then, ASMCs were immediately extracted by cutting the aorta along its length and by transferring the plane sample into a Phosphate Buffer Saline (PBS) bath. With tweezers, the adventitia was removed carefully in order to remove fibroblasts. The intima was removed as well and only the media was kept in the Petri dish. The media was cut into small pieces and immersed in tubes containing both elastase (Elastase, Lyophilized ESL, Worthington) and collagenase (Collagenase, Type I, powder, Gibco^TM^) in PBS. The tubes were heated to 37°C and shaken slowly for 3 hours until the final solution looked cloudy. In parallel, the culture flask was coated with fibronectin (Fn), using 10% Human Fn (Human Fn, Promocell) in PBS. The solution was kept 3 hours at room temperature or 30 minutes at the incubator before removing it from the flask. This coating was necessary for aortic SMCs in primary culture. Then, the solution was filtered successively into 70 µm and 40 µm strainers to eliminate the remaining ECM components and keep only the SMCs. The tube that contained the solution was carefully rinsed to filter three times with 10 mL PBS. After each filtration, the tube was centrifuged for 5 minutes at 1500 rpm, the supernatant was eliminated, and the pellet was suspended again with 10 mL PBS for the first time, and in 5 mL culture medium (SmGM-2, Lonza) at last. Finally, the cell suspension was transferred into the flask and completed with 5 mL of the medium. The flask was put into the incubator at 37°C and 5% CO_2_ for 2 weeks for sufficient cell growth, during two or three passages. At the end of this initial step of primary culture, the SMCs were frozen into 1.5 mL aliquots containing the same freezing solution as ASMC_1 and they were stored into liquid nitrogen. Each aliquot contained between 2 and 6 million SMCs.

##### 2.2.2 Cell culture and sample preparation

After thawing, the ASMC_1 and ASMC_2 cells were transferred into a T-75 flask for an entire week in the growth medium (SmGM-2, Lonza). The cells were incubated at 37°C and 5% CO_2_ to maintain the pH at 7.2-7.4. Then, ASMCs were cultured one week more in a basal medium (SmBM, Lonza), containing low (2%) FBS and 0.04% heparin, in order to preserve a contractile phenotype. Once they reached 50-70% confluence, a standard cell detachment protocol was used by applying a trypsin treatment with a low trypsin–EDTA solution (0.025% trypsin and 0.75 mM EDTA (1X), Sigma) to break down the focal adhesions in the culture dish without damaging the cells. Then, the cells in suspension could be used for subculturing or for sample preparation. ASMC_1 were seeded onto the sample surface at passage 5-6 and ASMC_2 at passage 3-4. Examples of ASMC_1 observed with fluorescence microscopy are shown in Figure 2.

Previously starved cells were transferred in three 24-well plates containing ready-to-use hydrogels with four different stiffness properties: 4, 8, 12 and 25 kPa. These hydrogels were made of a 400 µm-thick layer of polyacrylamide, which was assumed to be linear elastic within the range of strains considered in this study.^51^ The gel dimensions (12 mm diameter) were assumed to be infinitely large with respect to the cell size. Moreover, the collagen I coating added during the manufacturing process provided a physiological surface for cell adhesion and culture. About 10000 cells were seeded in each well and incubated in basal medium for two days before TFM experiments. This duration was sufficient to ensure spreading of SMCs, which adopted their specific elongated spindle shape.

##### 2.2.3 TFM measurements

A Carl Zeiss Axio Observer.Z1 station fitted in an incubating chamber was used to maintain the previously cultured hydrogels at 37°C and 5% CO_2_. According to the previously developed protocol,^32^ we recorded images of the gels at one frame per 30 seconds for a total duration of 5 minutes, during which SMCs were detached from the substrate by trypsin. Normally, cells detached from the gels within 1-2 minutes. We measured deformations induced by cell detachment in the gel by tracking the local motions of microbeads. For that, we magnified the area around several cells that showed a clear spindle shape. The field of view of the objective allowed the selection of 2-4 cells per well at the same time. The resulting images were processed using Digital Image Correlation (DIC) to obtain the corresponding displacement and strain fields around the focal adhesions of each ASMC (Figure 4c). Then a custom Matlab^®^ code of TFM analysis was applied for deriving the traction force values at each focal adhesion of interest.^32^

**Figure 4.**
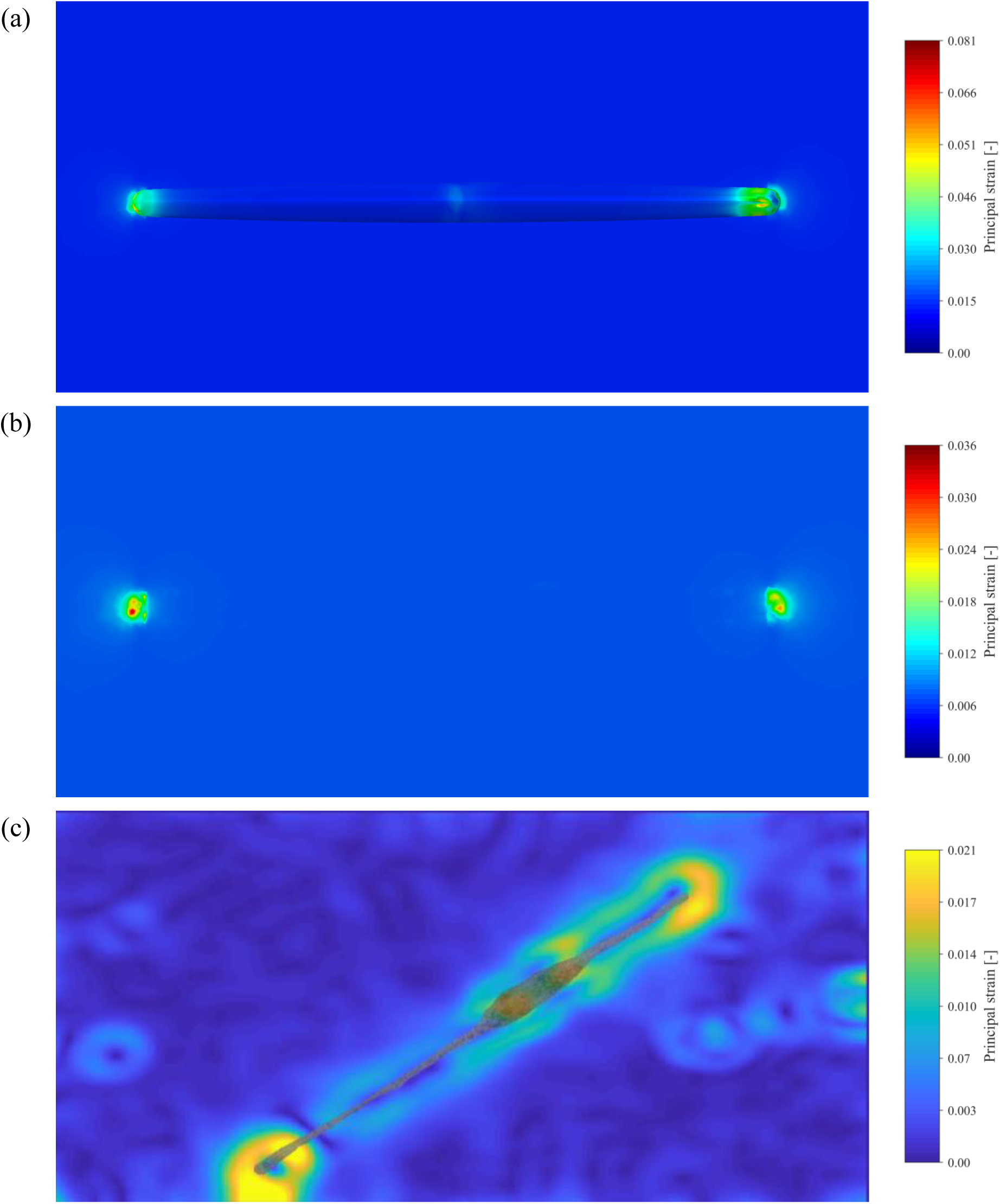
(a): Principal strain field simulated for a cell of the ASMC_1 lineage cultured on the 12 kPa substrate; the colormap of strain shown on the cell. (b): Principal train field simulated for a cell of the ASMC_1 lineage cultured on the 12 kPa substrate; colormap of strain shown on the surface of the substrate. (c): Strain field measured for a cell of the ASMC_1 lineage cultured on the 12 kPa substrate during the TFM experiments.

In total, traction forces were measured for 156 cells of the ASMC_1 lineage and for 169 cells of the ASMC_2 lineage. For the ASMC_1 lineage, 38 were cultured on gels of 4 kPa stiffness, 44 were cultured on gels of 8 kPa stiffness, 38 were cultured on gels of 12 kPa stiffness and 36 were cultured on gels of 25 kPa stiffness. For the ASMC_2 lineage, 37 were cultured on gels of 4 kPa stiffness, 50 were cultured on gels of 8 kPa stiffness, 44 were cultured on gels of 12 kPa stiffness and 38 were cultured on gels of 25 kPa stiffness. Results of TFM experiments were reported in details in another publication.^33^

#### 2.3 Parameter identification

To identify the constitutive parameters of our model introduced in subsection 2.1, we confronted its predictions with the TFM results obtained experimentally on the ASMC_1 and ASMC_2 cell lineages.

For each of the 8 groups of SMCs described in subsection 2.2, we analyzed the statistical distribution of the traction forces to derive the parameters of the Weibull probability density function of myosin activation. This needed first to normalize the experimentally measured traction forces (*TF*) such as:

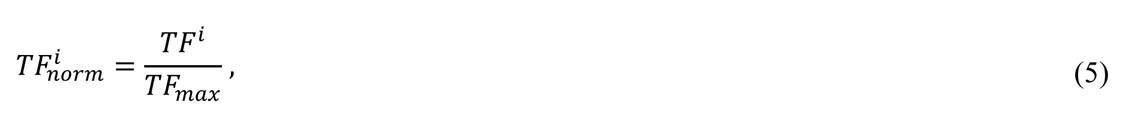

where *i* represents the sample number and *TF*_*max*_ is the maximum measured force. This normalization method was applied to each of the 8 datasets.

We derived the *k* and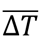 parameters of the Weibull functions best fitting the normalized traction force distributions for each of the 8 groups. Eventually, average values were deduced for each cell lineage (ASMC_1 and ASMC_2) and used to assign *k* and 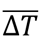 defining the myosin activation levels in the computational model.

Then, 300 cells were generated and simulated in the Abaqus software. Each cell had a random arrangement and a variable number of stress fibers and each activation level satisfied the previously defined Weibull distribution.

The contraction coefficient *α* was estimated for both ASMC_1 and ASMC_2 cell lineages by minimizing

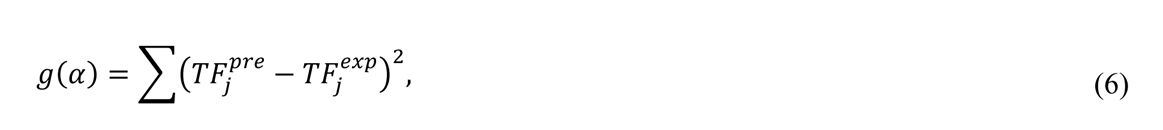

where *j* = {4, 8, 12}, and *TF*^j^_*pre*_ and *TF*^*exp*^_*j*_ represent the average predicted and the average measured traction forces, respectively, on substrates with elastic modulus 4, 8, and 12 kPa.

The elastic limit *σ*_*max*_ = *ξσ*_*SF*_*α*Δ*T* was set to the maximum stress value borne by stress fibers on the substrate with Young’s modulus 12 kPa, enabling the identification of *ξ*. Finally, *σ*^′^ was estimated by fitting the average predicted traction force on the stiffest substrate (25 kPa) to its corresponding experimental value.

## 3 Results

### 3.1 ASMC_1 cell lineage

For the ASMC_1 cell lineage, the distribution of traction forces measured experimentally showed an exponential shape (Figure 6). Therefore, the shape parameter was set to *k* = 1 in the Weibull probability density function. 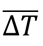 was estimated by fitting this exponential probability distribution function to the normalized traction forces measured for all the ASMC_1 cell lineage. Then, the contraction coefficient *α* was adjusted by fitting the model prediction obtained with Abaqus to the traction forces on substrates with the elastic modulus 4, 8, 12 kPa (25 kPa was excluded as it was used to estimate the softening part of the material behavior).

Maximum traction forces were measured on the 12 kPa substrate. Therefore, *σ*_*max*_ was set as the average stress value for this substrate and *ξ* was derived such as,

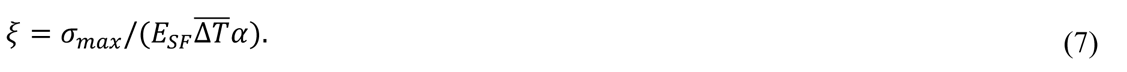

Lastly, *σ*^′^ was estimated by fitting the Abaqus results to the traction forces on the 25 kPa substrate. All the identified values for 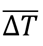, *α*, *ξ*, and *σ*^′^ are reported in Table 1.

**Table 1.**
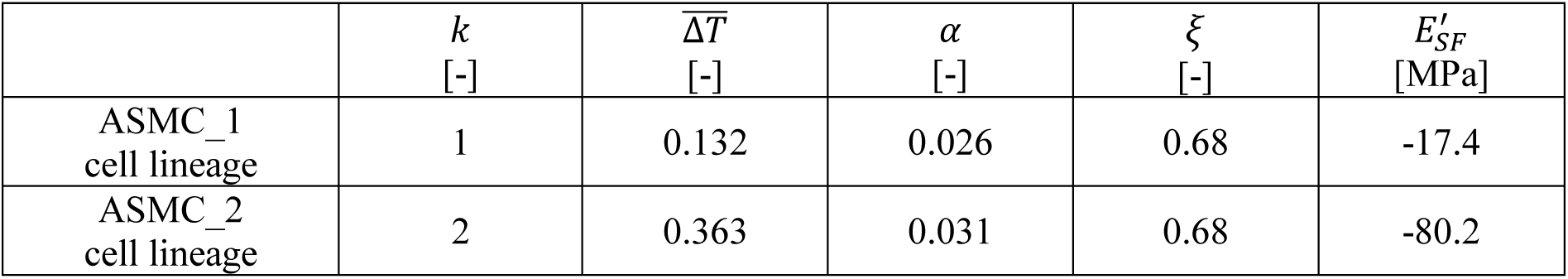
Model parameters estimated from the TFM experiments for the ASMC_1 and ASMC_2 cell lineages.

In Figure 4a, we show the strain field simulated for a cell of the ASMC_1 lineage with the FE model, using Δ*T* = 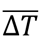 (basal tone) and 25 stress fibers, in which 13 of them connect one focal adhesion to the other one. In Figure 4b, we also show the simulated strain field on the substrate surface. The predicted traction force was *TF* = 31.7 nN for the 12 kPa stiffness substrate. Figure 4c also shows the measured absolute value for a cell cultured on 12 kPa substrate.

In Figure 5, we compare the measured traction forces with the average value predicted by the numerical model. To emphasize the role of stress fibers, two types of model predictions were plotted:

1. a first case (green curve) where we set *ξ* = 1 (equivalent to *σ*_*max*_= +∞), which corresponds to a fictitious cell with no softening effect in the stress-strain behavior of stress fibers.
2. a second case (blue curve), where the identified value *ξ* = 0.68 was assigned to the material model and where the softening behavior was driven by a *σ*^′^ modulus of - 17.4 MPa.

**Figure 5.**
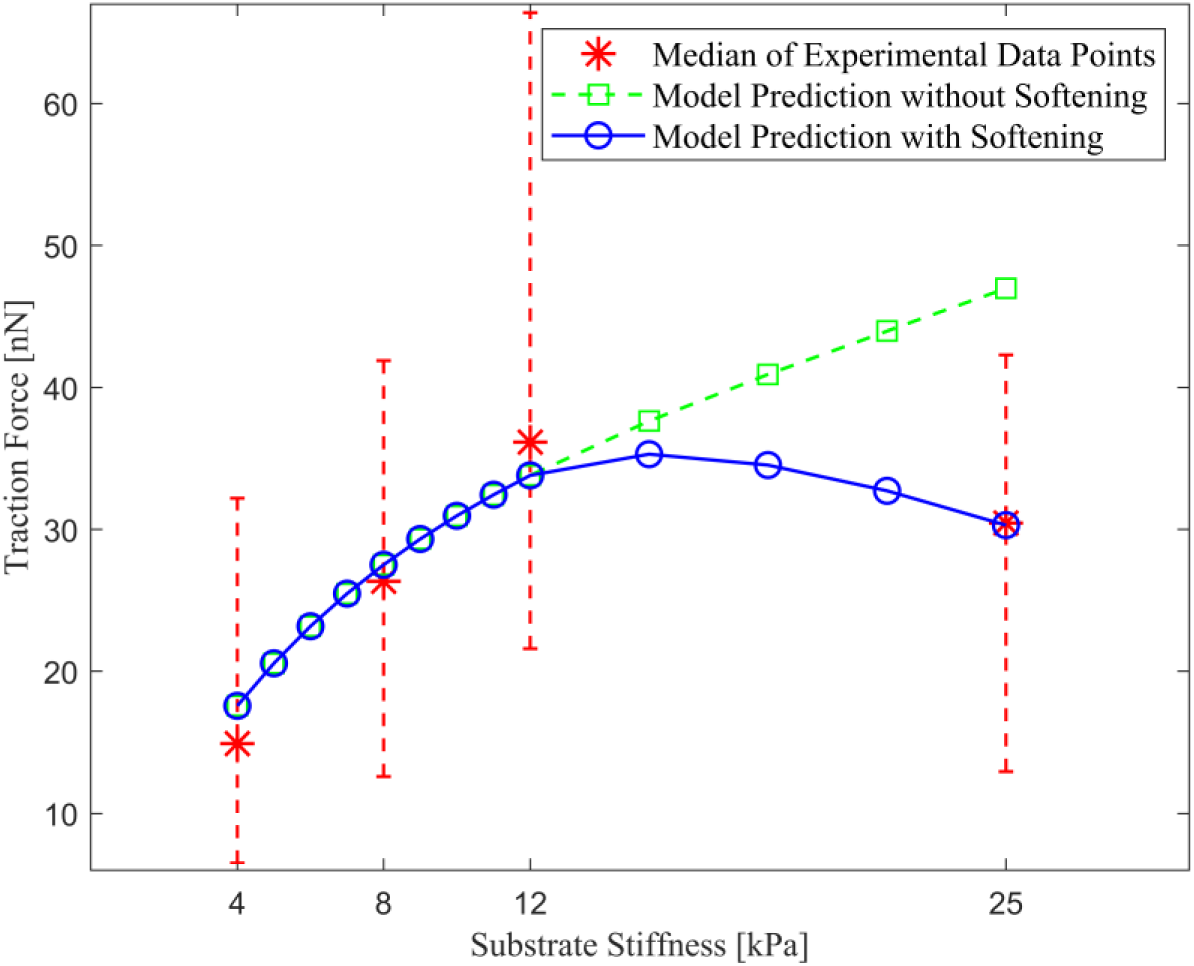
Comparison between the traction forces measured on the ASMC_1 lineage and their model predictions.

It can be observed that the identified model with the softening behavior is in very good agreement with the experimental results, whereas the absence of softening in the material model misses to reproduce precisely the stiffness sensitivity for substrates of large stiffness.

In Figure 6, we show the normalized histograms of contractile force values predicted by the numerical model superimposed with the normalized histograms of contractile force values measured on ASMC_1 cell populations. The statistical distribution predicted by the model was simulated for a virtual population of 300 SMCs with randomly varying numbers of stress fibers (following a uniform distribution) and randomly varying myosin activation levels Δ*T* (following the exponential distribution). There was a very good agreement between the simulated distribution and the model predictions. The remaining discrepancies were attributed to the dispersion of experimental data, which could probably be reduced by measuring traction forces on a larger population of cells.

**Figure 6.**
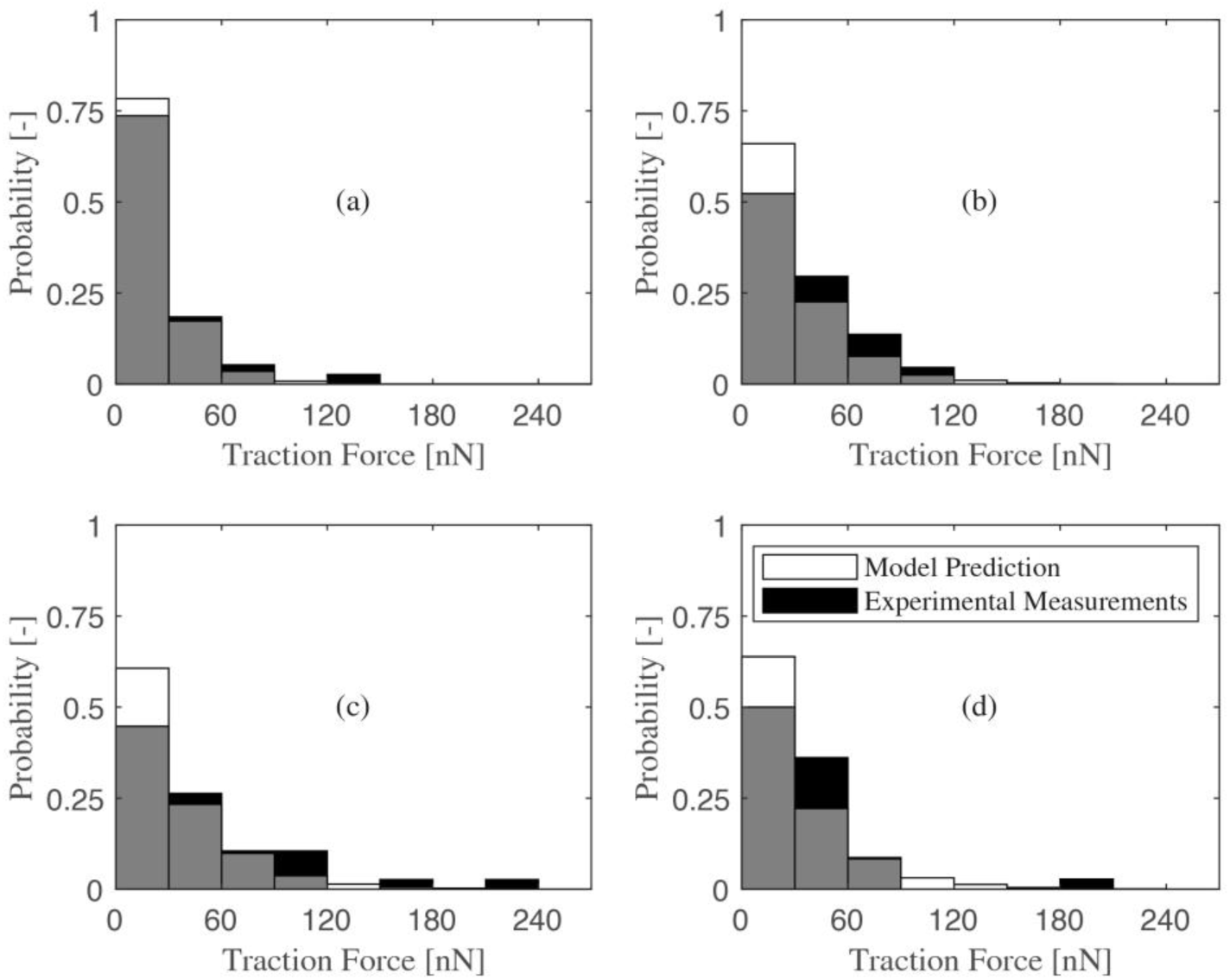
Histograms of the traction force distribution measured on the ASMC_1 lineage and predicted by the numerical model. The measured distributions show an exponential probability distribution function. (a): distribution of traction forces for cells cultured on the 4 kPa substrate; (b): distribution of traction forces for cells cultured on the 8 kPa substrate; (c): distribution of traction forces for cells cultured on the 12 kPa substrate; (d): distribution of traction forces for cells cultured on the 25 kPa substrate.

### 3.2 ASMC_2 cell lineage

A similar procedure was performed to estimate the model parameters for the ASMC_2 cell lineage. As the measured traction forces were significantly higher than for the ASMC_1 cells, a larger contraction coefficient *α* was estimated for the ASMC_2 cells. A sharper decrease of traction forces was also observed when SMCs were cultured on the stiffest substrate (25 kPa), resulting also in a larger value for the *σ*^′^ parameter.

Compared to the ASMC_1 cell lineage, the probability distribution of the normalized traction forces of the aneurysmal SMCs did not show an exponential form. It seemed to be similar to a Rayleigh probability distribution. Therefore, *k* = 2 was set in the defined Weibull probability density function. 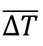 was estimated by fitting the Rayleigh probability distribution function to the normalized traction forces measured for all the ASMC_2 cell lineage. The identified model parameters are reported in Table 1.

In Figure 7, we compare the measured traction forces with the average value predicted by the numerical model for the ASMC_2 lineage. Again, it can be observed that the identified model with the softening behavior is in very good agreement with the experimental results, whereas the absence of softening in the material model would miss to reproduce precisely the stiffness sensitivity for substrates of large stiffness.

**Figure 7.**
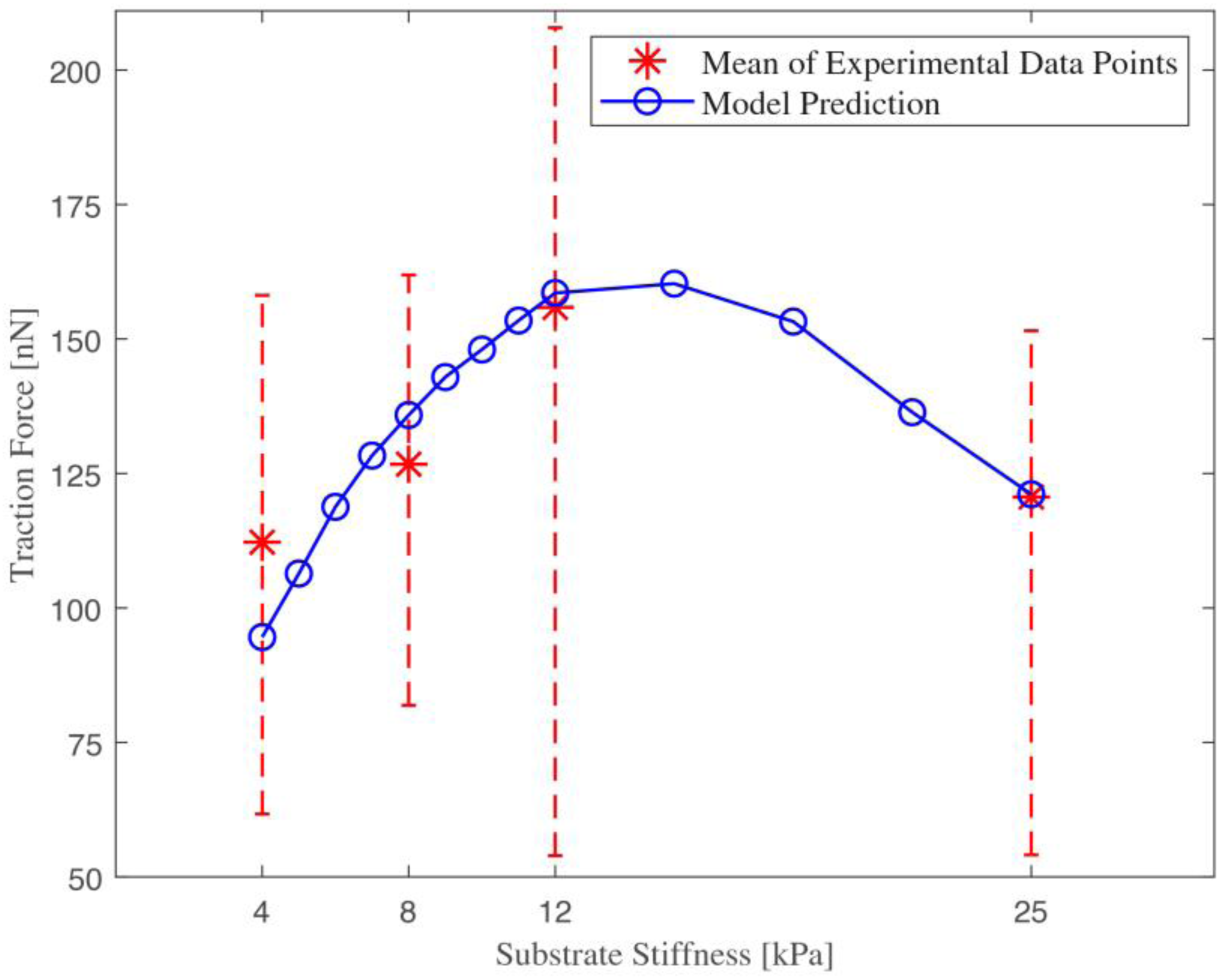
Comparison between the traction forces measured on the ASMC_2 lineage and their model predictions.

In Figure 8, we show the normalized histograms of contractile force values predicted by the numerical model superimposed with the normalized histograms of contractile force values measured on ASMC_2 cell populations. Similarly to the ASMC_1 cell lineage, a very good agreement was obtained between the simulated distribution and the model predictions. The remaining discrepancy could be attributed to the dispersion of experimental data, but also to some possible variations of 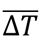 (median of myosin activation levels) between each group whereas the same 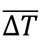 value (average of the 4 groups) was used for all the models.

**Figure 8.**
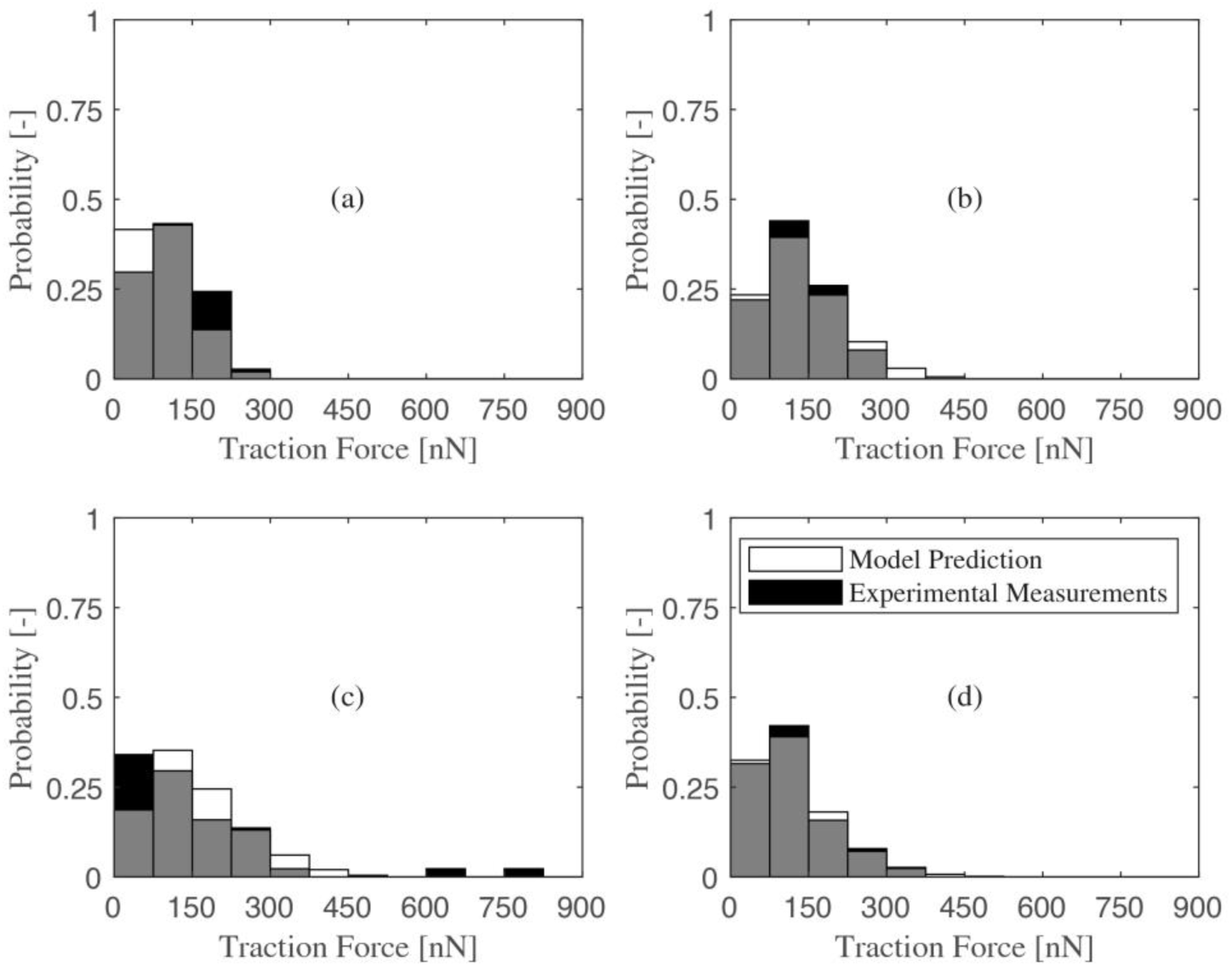
Histograms of the traction force distribution measured on the ASMC_2 lineage and predicted by the numerical model. The measured distributions show a Rayleigh-type probability distribution function. (a): distribution of traction forces for cells cultured on the 4 kPa substrate; (b): distribution of traction forces for cells cultured on the 8 kPa substrate; (c): distribution of traction forces for cells cultured on the 12 kPa substrate; (d): distribution of traction forces for cells cultured on the 25 kPa substrate.

### 3.3 Stress in the nuclear membrane and in the nucleus

The FE model was used to evaluate the stresses in the different cell components, especially the nuclear membrane and the nucleus. For each cell lineage and each substrate stiffness, a population of 300 cells were simulated and the average von Mises stress, as a scalar measure of the stress tensor, was deduced. Results are reported in Table 2. The stresses increase significantly when the substrate stiffness becomes larger. Unlike the traction forces at focal adhesions, the stresses in the nuclear envelope and in the nucleus keep increasing even for substrate stiffness beyond 12 kPa. Indeed, for substrates with larger stiffness, stress fibers connecting one focal adhesion to another one at both extremities of the cell reach their elastic limit, which causes their tension to drop. But stress fibers connecting one focal adhesion to the nuclear envelope can deform the cell nucleus instead of reaching the elastic limit. In the extreme case of a rigid substrate, the strain of stress fibers *ε*_*a*_ introduced in Equation (3) is null for stress fibers attached to the substrate at both extremities, whereas it can be approximated such as *ε*_*a*_ = *ε*_*nucl*_ when one extremity is attached to the nuclear envelope, where *ε*_*nucl*_ is the strain resulting from elastic deformation of the nucleus. Therefore, the tensile stress of a stress fiber attached to the nuclear envelope would write:

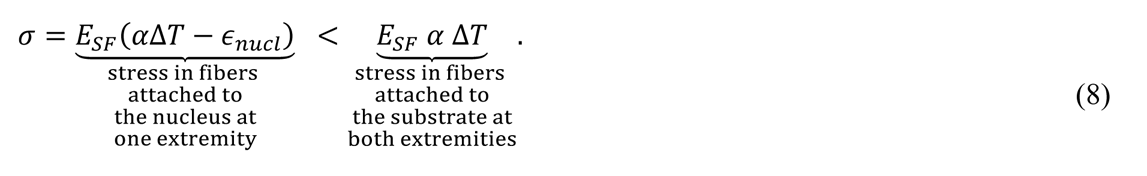

**Table 2.** Average von Mises stress values in the nucleus and nuclear membrane computed for ASMC_1 and ASMC_2 with the computational model.

| Average von Mises stress |  | Substrate Stiffness [kPa] |  |  |  |
| --- | --- | --- | --- | --- | --- |
|  |  | 4 | 8 | 12 | 25 |
| ASMC_1 | Nucleus [Pa] | 0.665 | 0.844 | 0.984 | 1.187 |
|  | Nuclear Membrane [kPa] | 17.039 | 21.578 | 24.933 | 30.163 |
| ASMC_2 | Nucleus [Pa] | 2.197 | 2.789 | 3.251 | 3.994 |
|  | Nuclear Membrane [kPa] | 56.292 | 71.288 | 82.371 | 101.369 |

Moreover, we studied the sensitivity of mechanical properties of the nuclear membrane on the traction forces and average von Mises stress values. For the substrate with the elastic modulus of 12 kPa, figure 9a shows the changes in the traction force for ASMC_1. This figure shows ratios of the average traction forces among 300 different cells to the reference case in which the shear modulus of the nuclear membrane is 600 kPa. The results show that as the nuclear membrane becomes stiffer, the stress fibers connecting one focal adhesion to the nuclear envelope are more constrained and have more contribution to the cell force-generating capacity. However, the changes of traction forces are not significant. It can be explained by the fact that the average number of stress fibers connecting one focal adhesion to the other one was 19, and the average number of fibers connecting one focal adhesion to the nuclear envelope was 6. Therefore, the force-generating capacity mainly depended on the stress fibers connecting focal adhesions to each other. Besides, figure 9b shows the changes in the average von Mises stress values in the nucleus and in the nuclear membrane. For a more compliant nuclear membrane, the average von Mises stress reduces in the nuclear envelope but in the nucleus, as it undergoes higher strain values, we observe an increased average von Mises stress.

**Figure 9.**
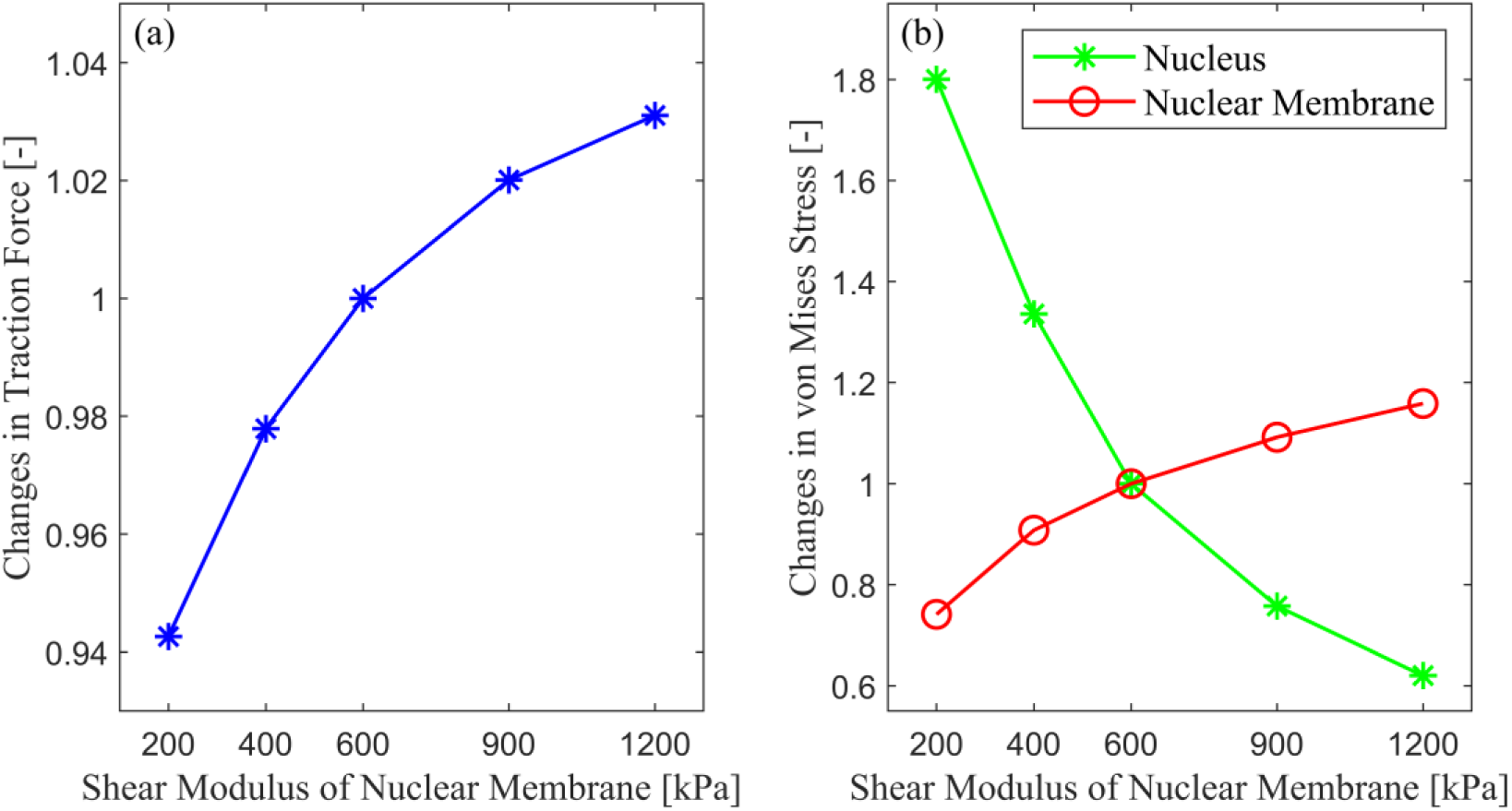
Sensitivity of the mechanical properties of the nuclear membrane on the traction force and average von Mises stresses on the 12 kPa substrate. (a): ratios of the traction force to the case in which the shear modulus of the nuclear membrane is 600 kPa. (b): ratios of the average von Mises stress values in the nucleus and nuclear membrane to the case in which the shear modulus of the nuclear membrane is 600 kPa.

### 3.4 Effects of SMC width

Lastly, we investigated the effects of the width of SMCs on their force-generating capacity. As shown in Figure 2, the width of ASMCs can vary significantly from one cell to another. To compare the results with the main model shown in figure 3a, we repeated the FE simulations for 2 other cases. For the first one, a scale factor of 0.5 was applied to the width of the cell and to the nucleus radios, and for the second one, a scale factor of 2 was applied. Moreover, we kept the same density of stress fibers for all cases. Figure 10 shows the obtained traction forces for different widths. It shows that the traction force linearly increases with the width increase and therefore with the number of stress fibers. Moreover, Figure 10b shows the changes in the average von Mises stress values in both the nucleus and nuclear membrane. By increasing the width, the striking result is that the nucleus undergoes higher stresses compared to the nuclear membrane.

**Figure 10.**
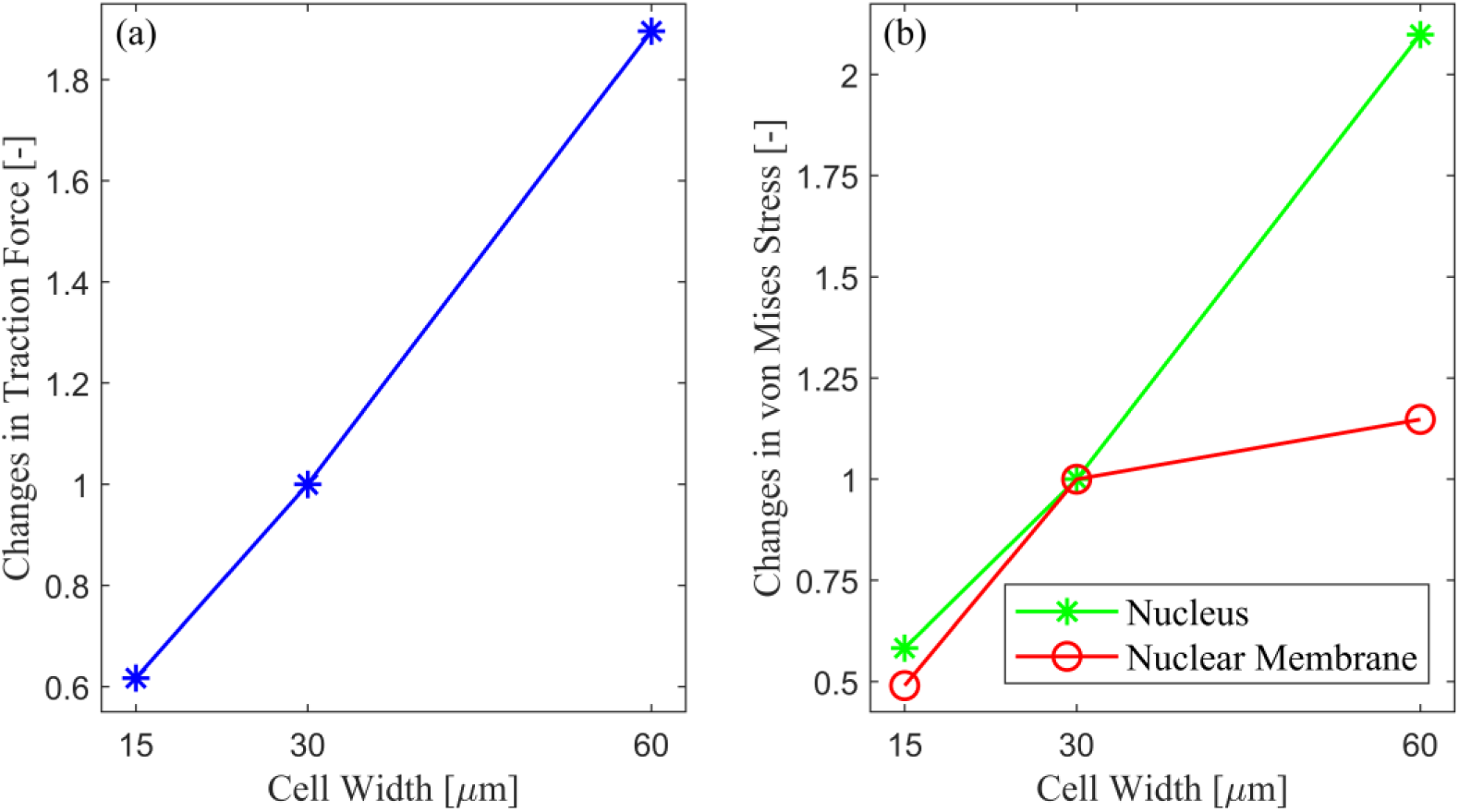
Effects of the width of SMCs on their force-generating capacity (a) and stress values in the nucleus and nuclear membrane (b). All cases were compared to the reference case in which the width is 30 μm.

## 4. Discussion

Dysfunction in the ability of SMCs to sense mechanical stiffness and respond to its changes has been shown to be a common factor in many vascular diseases.^2, 22, 24^ However, the mechanisms that underlie the ability of SMCs to sense and respond to stiffness change are still partially unclear. Several previous studies on different cell types have examined the roles of mechanoreceptors at the cellular membrane.^25–27^ Moreover, recent studies have highlighted the significant role of contractile acto-myosin units in the cytoskeleton that can act as stiffness sensors.^28–30^ In the current study, we have developed the first continuum mechanics model of SMCs accounting for stiffness sensing through the contractile acto-myosin units.

Cells are known to modulate their traction force in response to changes in substrate stiffness.^52^ For a given cell type cultured on an elastic substrate, traction forces can be divided into two regimes. In the first regime, for stiffness values below a threshold, the cell develops a traction force which is related to the substrate stiffness. Beyond this threshold, however, the traction force exerted by the cell remains largely insensitive to further changes in substrate stiffness.^28^ Our experimental results confirmed that ASMCs do modulate their traction forces according to these two regimes. We even observed a decrease of the traction forces in the second regime beyond the stiffness threshold. The stiffness threshold was found at 12 kPa. This is consistent with other studies which found that the striation of myotubes was maximum for substrate rigidities around 12 kPa.^28^ This is also consistent with predictions of the motor–clutch-based model, which show that there exist an ‘optimal stiffness’ where cells can generate maximal traction.^46^

The main goal of our study was to provide a simple continuum mechanics model of ASMCs accounting for stiffness sensing. This was achieved by modeling stress fibers as truss-like structures embedded in a continuum medium representing the cytoplasm. Within a living cell, myosin motor proteins use the energy from ATP hydrolysis to repetitively bind to actin and walk along the actin fiber toward its positive end. When the load on the myosin crosslink approaches the stall force, the myosin takes backward and forward steps around the same location.^29^ Moreover, recent findings on cells showed that in the rigidity sensing process, the myosin filaments contract the actin filaments to a fixed distance.^44^ Inspired by this, we considered a strain-driven behavior for the contraction of stress fibers, which was introduced by the eigenstrian *α*Δ*T* in section 2.1. In our model, these effects involved in the contraction of stress fibers have been simply modeled with the concept of analogous thermal strains. Without any complex equations, the proposed model can simulate stress fiber contraction and fit the actual traction forces reported by TFM experiments. The sliding effects when the myosin crosslink approaches the stall force have been transduced into a softening constitutive behavior of the truss-like structure, with an elastic limit *σ*_*max*_ and a softening coefficient *σ*^′^.

The cellular environment can only affect the regulation of essential cellular processes (DNA replication, chromatin organization, cell division, differentiation) if a signal is sensed by the cell nucleus. Nuclear mechanosensing was modeled by considering a perinuclear actin cap connecting a fraction of stress fibers to the interphase nucleus through LINC protein complexes.^36, 37^ Recently, Nagayama obtained a direct evidence for the mechanical interaction between stress fibers and the nucleus by using a laser-based nano-dissection technique.^53^ Briefly, they cut a stress fiber running across the top surface of the nucleus by using a laser to release its pretension, and observed the resultant deformation of the dissected stress fiber and the nucleus. The findings indicate that this mechanical interaction may achieve direct force from stress fiber to the nucleus and induce conformational changes in the intranuclear chromatin.

Our model shows that the substrate stiffness directly affects the tension in the nuclear envelope (actin perinuclear cap) and the stress in the nucleus. Accordingly, forces transferred via the cytoskeleton can directly alter gene expressions by inducing deformations of the nucleus, as accumulating evidence suggests that the three-dimensional organization of the nucleus regulates gene expression through laminae chromosome interactions.

Forces transferred via the cytoskeleton can also affect the posttranscriptional control of gene expression by causing nuclear pores to open.^54^ Indeed, deformations of the nuclear envelope and of the actin perinuclear cap may be required for universal mechanotransducers YAP/TAZ to translocate into the nucleus.^37, 55^

Our model only accounts for stiffness sensitivity through the acto-myosin contractility mechanisms. In reality, these mechanisms may act synergistically with the mechanosensitivity of focal contacts.^28^ For instance, myosin-dependent sensitivity may adapt cytoskeletal tension to substrate rigidity at the scale of the whole cell, this tension being then transduced locally through focal contacts sensitivity. Integrins have been shown to act as mechanosensors which in turn interacts with other signaling molecules to trigger many downstream signaling cascades.^26, 56^ Mechanical forces can also directly cause the activation of ion channels and be sensed by the nucleus through the cytoskeleton. Biochemical signaling also plays a critical role in the response of the cell to stretch and substrate stiffness. Indeed, the mechanisms based on the actin-myosin network presented here are only one part of a complex mechanosensing and response machinery in the cell and act in conjunction with these already established molecular signaling pathways that have important mechanosensory roles.^56^

Another important aspect of our computational model is to account for the intercellular variability of acto-myosin contractility. The variability is modeled through a random arrangement of stress fibers, a variable number of stress fibers, and a distribution of activation levels satisfying the Weibull probability density function. Stress fibers, as the major contributor to the contractile behavior, are randomly distributed inside the cells.^57^

Most importantly, the predictions of our computational model were in agreement with experimental measurements of traction forces. Petit and colleagues found that the larger traction forces in the aneurysmal SMCs compared to the healthy ones are related to the larger size and higher density of stress fibers in the aneurysmal SMCs.^33^ We showed that our model can predict this alteration in force-generating capacity.

To the authors’ knowledge, the proposed model is the first continuum mechanics model that explains the stiffness sensitivity of cells by the mechanical properties of cells. Although it can provide an accurate prediction for the contractile force capacity of both healthy and aneurysmal SMCs, it still suffers from a number of limitations:

1. We modeled only the effects of the basal tone of SMCs through a quasi-static approach. We disregarded all the dynamics of signal transmission through stress fibers,^58^ dynamics of the actin-myosin contractility process, dynamics of actin polymerization, or model coordination with cell-specific processes like spreading and cytoskeletal remodeling. It is known that the active motion of myosin motors within such a network can also result in rearrangement of the actin network.^29^ These dynamical effects will be integrated in further developments of the model, which will include Huxley’s sliding filaments theory accounting for the effect of the acto-myosin detachment on the velocity of contractile shortening.^43^
2. Although the proposed model predictions fairly follow the experimentally obtained histogram, there are some slight differences between experimental results and model predictions. One of the reasons for this discrepancy may be the relatively low number of samples. However, the main trends of stiffness sensing expressed in the experiments by ASMCs were caught correctly by the model.
3. Stress fibers were modeled with truss-like elements. These elements cannot bear compressive loads which are transferred to the surrounding cytoplasm. Further developments of the model should include the role of microtubules which have also been shown to contribute to cell mechanosensitivity.^59^
4. Although experimental investigation on single stress fibers has shown a nonlinear stress-strain response,^60^ for the sake of simplicity, stress fibers were assumed to behave linear elastically in our model. Moreover, the material model of all the other components of the cell was simply a Neo-Hookean model, with material parameters obtained from the literature. Nevertheless, more sophisticated material models can be easily used in the future provided that they can be calibrated with appropriate experimental characterizations.

## 5. Conclusion

Any dysfunction in SMCs contractility can affect the mechanical response and load-bearing capacity of arteries. This study was dedicated to developing a new mechanical model to simulate stiffness sensing by SMCs. We focused on the role of acto-myosin contractility in stiffness sensing and introduced a universal stress-strain relationship for stress fibers. Model predictions were in very good agreement with cellular tractions measured on different cell lineages. Finally, stresses in the nuclear envelope and in the nucleus were computed with the model, showing that the variations of cytoskeletal forces induced by substrate stiffness directly induce deformations of the nucleus which can potentially alter gene expression.

The predictability of the model combined to its relative simplicity are promising assets for further investigation of stiffness sensing in 3D environments. Eventually, this could contribute to decipher the effects of mechanosensitivity impairment, which are known to be at the root of aortic aneurysms.^2, 23^ Extensions of the model to other types of adherent cell such as fibroblasts or endothelial cells is also envisaged in the future.^47^

## Acknowledgements

The authors are grateful to the ERC for financial support through ERC-2014-CoG BIOLOCHANICS grant.

## Author declarations Conflict of Interest

The authors declare that they have no conflict of interest related to the present study.

## Ethics Approval

Primary cells were obtained with the authorization of the French Biomedicine Agency—PFS 09–007—in accordance with the declaration of Helsinki) and, after informed consent, on a patient undergoing ATAA surgical repair according to protocols approved by the CHU-SE ethics committee (Centre Hospital- Universitaire ids—Saint-Etienne, France).

## Author Contributions

**Ali Akbar Karkhaneh Yousefi**: Conceptualization (equal); Methodology (lead); Software (lead); Writing – original draft (lead); Writing – review & editing (equal). **Claudie Petit**: Writing – original draft (supporting); Resources (equal); Writing – review & editing (equal). **Amira Bin Hassine**: Resources (equal); Writing – review & editing (equal). **Stéphane Avril**: Conceptualization (equal); Funding acquisition (lead); Supervision (lead); Writing – review & editing (equal).

## Notes

### Competing Interest Statement

The authors have declared no competing interest.

